# FALDO: A semantic standard for describing the location of nucleotide and protein feature annotation

**DOI:** 10.1101/002121

**Authors:** Jerven Bolleman, Christopher J. Mungall, Francesco Strozzi, Joachim Baran, Michel Dumontier, Raoul J. P. Bonnal, Robert Buels, Robert Hoehndorf, Takatomo Fujisawa, Toshi-aki Katayama, Peter J. A. Cock

## Abstract

**Background** Nucleotide and protein sequence feature annotations are essential to understand biology on the genomic, transcriptomic, and proteomic level. Using Semantic Web technologies to query biological annotations, there was no standard that described this potentially complex location information as subject-predicate-object triples.

**Description** We have developed an ontology, the Feature Annotation Location Description Ontology (FALDO), to describe the positions of annotated features on linear and circular sequences. FALDO can be used to describe nucleotide features in sequence records, protein annotations, and glycan binding sites, among other features in coordinate systems of the aforementioned “omics” areas. Using the same data format to represent sequence positions that are independent of file formats allows us to integrate sequence data from multiple sources and data types. The genome browser JBrowse is used to demonstrate accessing multiple SPARQL endpoints to display genomic feature annotations, as well as protein annotations from UniProt mapped to genomic locations.

**Conclusions** Our ontology allows users to uniformly describe – and potentially merge – sequence annotations from multiple sources. Data sources using FALDO can prospectively be retrieved using federalised SPARQL queries against public SPARQL endpoints and/or local private triple stores.

## Background

Describing regions of biological sequences is a vital part of genome and protein sequence annotation, and in areas beyond this such as describing modifications related to DNA methylation or glycosylation of proteins. Such regions range from one amino acid (e.g. phosphorylation sites in singalling cascades) to multi megabase contigs mapped to a complete genome. Such annotation has been discussed in biological literature since at least since 1949 [1] and recorded in biological databases since the first issue of the Atlas of Protein Sequence and Structure [2] in 1965.

There are many different conventions for storing genomic data and its annotations in plain text flat file formats such as GFF3, GVF [3], GTF and VCF, and more structured domain specific formats such as those from INSDC or UniProt, but none are flexible enough to discuss all aspects of genetics or proteomics. Furthermore, the fundamental designs of these formats are inconsistent, for example both zero-based and one-based counting standards exist, a regular source of off-by-one programming errors which experienced bioinformaticians learn to look out for.

Although non-trivial, file format interconversion is a common background task in current script-centric bioinformatics pipelines, often essential for combining tools supporting different formats or format variants. As a result of this common need, file format parsing is a particular strength of community developed open source bioinformatics libraries like BioPerl [4], Biopython [5], BioRuby [6] and BioJava [7]. While using such shared libraries can reduce the programmer time spent dealing with different file formats, adopting Semantic Web technologies has even greater potential to simplify data integration tasks.

As part of the Integrated Database Project (http://lifesciencedb.mext.go.jp/en/) and the Core Technology Development Program (http://biosciencedbc.jp/en/tec-dev-prog/programs) to integrate life science databases in Japan, the National Bioscience Database Center (NBDC) and the Database Center for Life Science (DBCLS) have hosted an annual “BioHackathon” series of meetings bringing together biological database teams, open source programmers, and domain experts in Semantic Web and Linked Data [8–11]. At these meetings it was recognised that failure to standardise how to describe positions and regions on biological sequences would be an obstacle to the adoption of federalised SPARQL/RDF queries which have the potential to enable cross-database queries and analyses. Discussion and prototyping with representatives from major sequence databases such as UniProt [12], DDBJ [13] (part of the INSDC partnership with the NCBI-GenBank [14] and EMBL-Bank [15]), and a number of glycomics databases (BCSDB [16], GlycomeDB [17], GLYCOSCIENCES.de [18], JCGGDB, RINGS [19] and UniCarbKB [20]) and assorted open source developers during these meetings led to the development of the Feature Annotation Location Description Ontology (FALDO).

FALDO has been designed to be general enough to describe the position of annotations on nucleotide and protein sequences using the various levels of location complexity used in major databases such as INSDC (DDBJ, NCBI-GenBank and EMBL-Bank) and UniProt, their associated file formats, and other generic annotation file formats such as BED, GTF and GFF3. It includes compound locations, which are the combination of several regions (such as the ‘join’ location string in INSDC), as well as ambiguous positions. It allows us to accurately describe ambiguous positions today in such a way that future more precise knowledge does not introduce logical conflicts which potentially could only be resolved by intervention of an expert in the field.

FALDO is suited to accurately describe the position of a feature on multiple sequences. This is expected to be most useful when lifting annotation from one draft assembly version to another. For example, a gene can start at a position for a given species’ genome assembly, while the conceptually same gene can start at another position in previous/following genome assemblies for the species in question.

FALDO has a deliberately narrow scope which does not address general annotation issues about the meaning of or evidence for a location, rather FALDO is intended be used in combination with other relevant ontologies such as the Sequence Ontology (SO) [21] or database-specific ontologies. That is, it is used only to describe the loci of features, not to describe the features themselves. A FALDO position relative to a sequence record is comparable to a coordinate position on a map: it makes no claim about how that sequence record or map is related to the real world.

## Implementation

FALDO is a small OWL2 ontology with 14 classes of which 9 deal with the concept of a position on a sequence (Figure 1). Four of those classes are used to describe accurately what we know of a position that is not precisely determined. Four classes are used to describe the concept of a position on a strand of DNA, e.g. positive, negative and on both strands. All eight of these classes are sub classes of the generic faldo: Position super-class. The ninth class is the concept of a region i.e. something with a end and start position. In contrast to other representations, FALDO has no explicit way to say that it is not “known” on which strand a position is, because this explicit statement unknown strand position can introduce contradictions when merging different data sets. For example, some positions could end up being contradictorily typed both as forward-stranded as well as being located on an unknown strand position.

**Figure 1:**
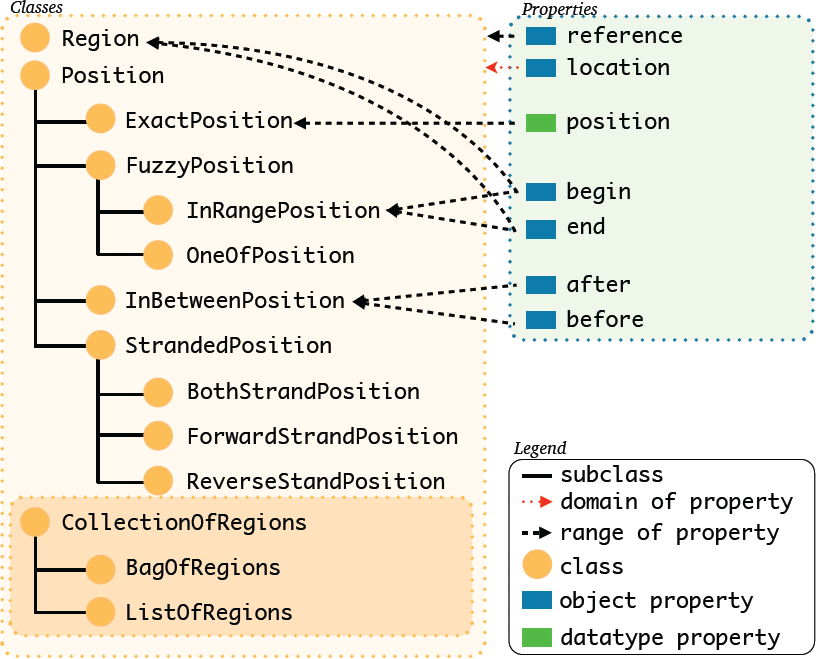
The classes and object properties used in FALDO.

**Figure 2:**
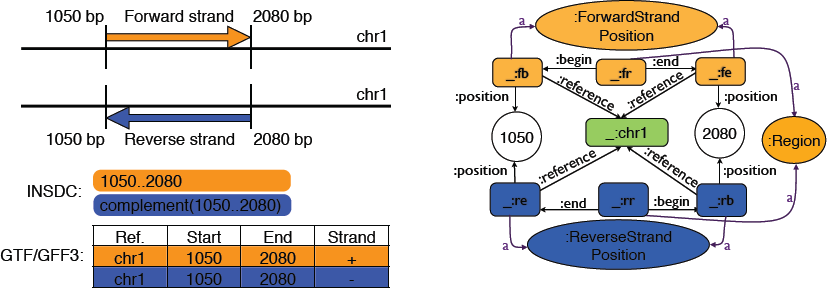
Assorted conventions for regions, start, end, and strands. This figure shows two hypothetical features on a DNA sequence (labeled chr1), on either the forward strand (orange) or reverse strand (blue). Using the INSDC location string notation, these regions are “1050..2080” and “complement(1050..2080)” respectively if implicitly given in terms of the reference chr1. Using the GTF/GFF3 family of formats, regardless of the strand these two locations are described with *start* = 1050 and *end* = 2080, and in general, *start ≤ end*. Biologically speaking, in terms of transcription, the start of a genomic feature is strand dependent. For the forward strand feature (orange), the start is 1050 while the reverse strand feature (blue) starts from 2080.

There are 3 more classes (faldo: CollectionOfRegions and its subclasses) that are only there for backwards compatibility with INSDC join features with uncertain semantics. i.e. those join regions where a conversion program can only state that there are some regions and that the order that they are declared in the INSDC record might have biological significance. However, here the INSDC record needs intelligent inspection before the data can be cleanly converted to a data model with rich semantics.

FALDO defines a single datatype property, faldo: position, that is used to provide a one-based integer offset from the start of a reference sequence. This property, when used together with the faldo: reference property, links the concept of a faldo: Position to an instance of a biological sequence. Note that these terms are case-sensitive: faldo: position is a property, and faldo: Position is a concept.

For compatibility with a wide range of data, FALDO makes very few assumptions about the representation of the reference sequence, and can be used to describe positions on both single- and double-stranded sequences. When both strands of a double-stranded sequence are represented by a single entity (recommended over each strand being represented separately), integer faldo: position properties are counted from the 5’ end of whichever strand is considered the “forward” strand.

A key part of the FALDO model is the separation of feature and where a feature is found in a sequence record. For this we use the faldo: location object property. This property is used to distinguish between a conceptual gene as an “unit of inheritance” and the corresponding representation of the DNA sequence region encoding the gene as stored in a database.

As in the INSDC data model and the associated GenBank ASN.1 notation, each location in FALDO has an identifier for the sequence it is found on [22]. This means that the position information is complete without further references to the context the position information was found in. The difference is that in FALDO, due to its RDF nature, the identifier of the sequence is a dereferencable pointer (URI) on the web, instead of just a string of characters.

### Compression via OWL2 reasoning

For large databases such as INSDC or UniProt, the need to repeat the reference sequence for each position may come with a significant cost in storage. However, this triple does not need to be materialised in the database, as it is inferrable using OWL2 property chain reasoning. With the axiom shown in Figure 3 the faldo: reference triples can be inferred for any faldo: position described by an INSDC record. Having an OWL-capable query rewriter allows users to ignore the difference between encoding the faldo: reference properties explicitly and having them inferred at query time. For RDF databases that do not offer this capability, the necessary triples can be easily added using a single SPARQL insert query (Figure 4). This flexibility allows users of the data to select the best approach for their infrastructure, rather than being constrained by the decisions of the data provider.

**Figure 3:**
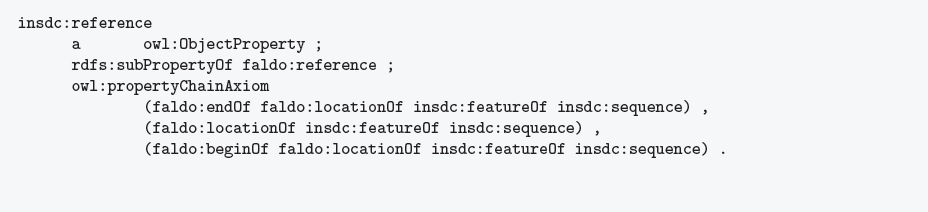
OWL2 property chain axiom to infer that all positions described in an INSDC record are relative to the main sequence of the record (in RDF turtle syntax, prefixes ommited).

**Figure 4:**
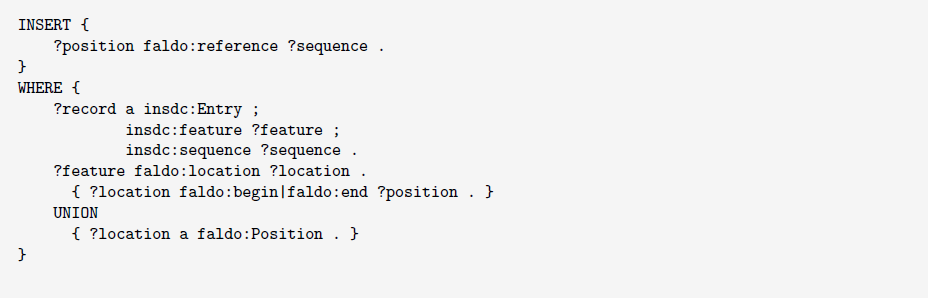
A SPARQL query to add all faldo: reference properties to faldo: positions described from a insdc: record.

### Validating data encoded with FALDO

Some databases only allow a subset of FALDO. For example INSDC requires that the start and end of a region are on the same sequence, while UniProt requires that a feature is described in relation to the reference’s canonical isoform. Yet another database might annotate the location of a glycsoylation site on an UniProt isoform sequence. When added to an UniProt record in RDF, this extra RDF annotation would be ignored by applications that are not concerned with glycosylation of isoforms. The same annotation can not be added to UniProt XML as the XSD schema does not allow for it, and the older plain text flat-file format does not allow for this kind of third party extension either. An attempt to add such information would very likely break any XML or flat-file parser and introduces the risk of importing data incorrectly. Only the UniProt RDF format allows other people to make assertions about UniProt data without breaking existing tools.

There are many ways to add constraints to the data model by applications using Semantic Web technologies [23]. In other words, data validation is an application specific concern instead of a data format concern.

### Users

FALDO is already deployed and used in a number of tools and databases.

**JBrowse** can use SPARQL queries with FALDO to visualize annotations on reference sequences from semantic databases [24] (see Figure 5).
**INSDC-DDBJ** DDBJ is currently working on an RDF format for the INSDC data that is stored in DDBJ/GenBank/EMBL-Bank.
**BioInterchange** uses FALDO to make position information stored in current bioinformatics formats (s.a. GFF3, GTF and GVF) available to the Semantic Web (http://www.biointerchange.org/).
**TogoGenome** a genome database collection provided by the DBCLS also uses FALDO in its RDF representation (http://togogenome.org/).
**PhenomeBrowser** The positions on the mouse genome of phenotype and disease related natural variations are described using FALDO.
**BOING** The “bio-ontology integrated querying of sequence annotations” framework uses FALDO to describe all feature locations [25].
**SPARQL-BED** This simple tool that turns any BED file into a Web accessible SPARQL endpoint using FALDO to describe BED feature positions (https://github.com/JervenBolleman/sparql-bed).
**BioPerl** BioPerl [4] now includes a FALDO exporter (Bio::FeatureIO::faldo), which allows any BioPerl-supported feature format to be translated to FALDO.
**UniProt** UniProt annotates many protein features and sites. Starting with UniProt RDF release 2014 01 the positions of protein feature are described using FALDO.

**Figure 5:**
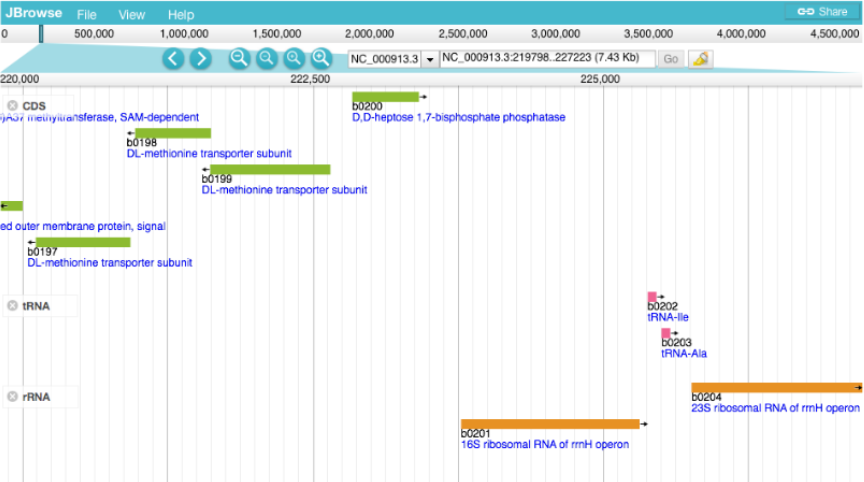
JBrowse showing features, whose location is encoded using FALDO, selected via SPARQL (at http://togogenome.org/).

## Results

One of the practical goals driving the development of FALDO was to be able to represent all the annotated sequences in INSDC and UniProt as RDF triples, as a step towards providing this data via SPARQL endpoints where it can be queried.

The protein examples considered here, such as the UniProt feature annotations, describe relatively simple locations within protein sequences (see the active site annotation in Figures 6 and 7).

**Figure 6:**
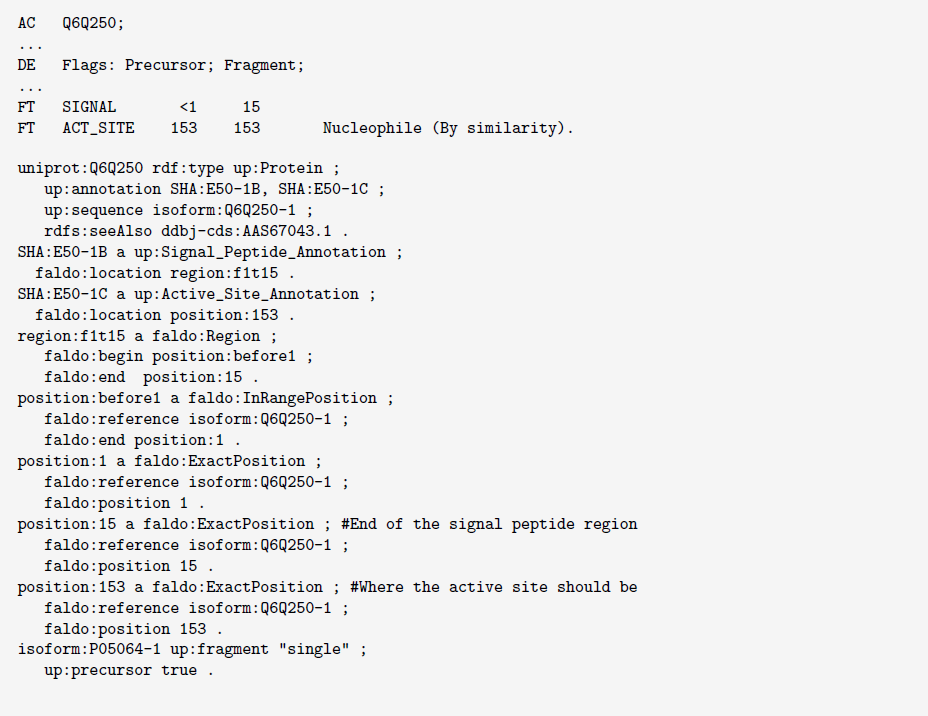
Excerpt from UniProt entry Q6Q250 showing the position of an active site and a signal peptide in both the UniProt flat-file format and FALDO.

**Figure 7:**
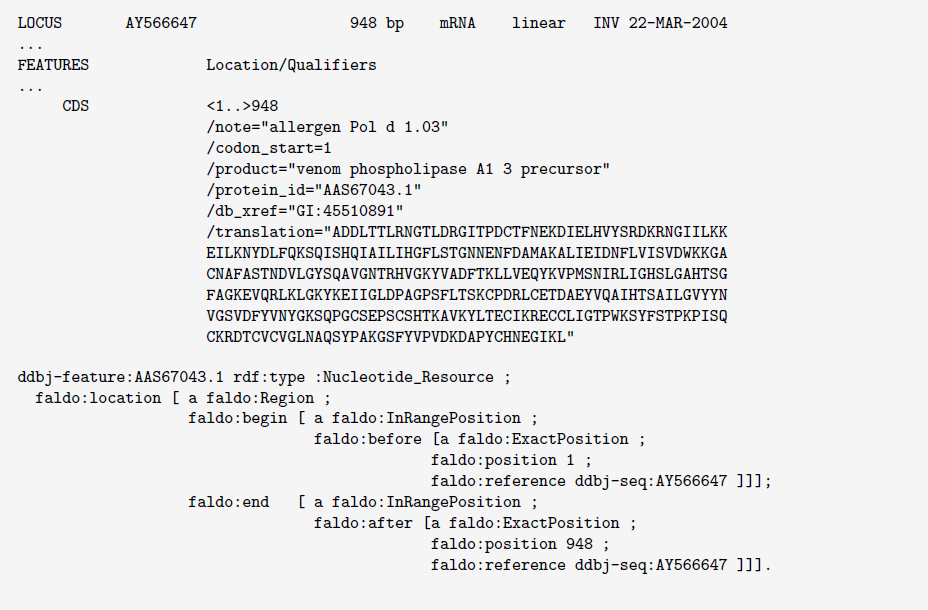
DDBJ record associated with UniProt Q6Q250 showing the related CDS sequence, with coding region outside of the known deposited mRNA sequence.

### Complement strand and INSDC compound locations

Describing biological features in relation to a genomic DNA sequence does not have to be complicated.

For example the *cheY* gene (shown in Figure 8) *Escherichia coli* str. K-12 substr. MG1655 (accession NC 000913.2) is described in the INSDC feature table as complement(1965072..1965461), which is 390 base pairs using inclusive one-based counting. This feature begins on the base complementary to *start* = 1965461 and finishes at *end* = 1965072, so the INSDC location string can be interpreted as complement(*end*..*start*). FALDO respects this biological interpretation of a feature location on the reverse strand.

**Figure 8:**
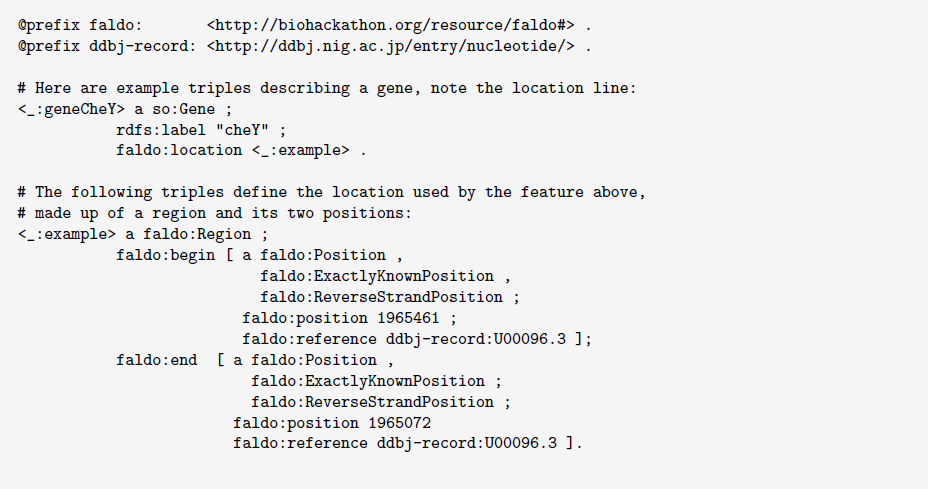
Using FALDO in Turtle [26] syntax to describe the location of a gene feature *cheY* at complement(NC 000913.2:1965072..1965461) in a INSDC record U00096.3.

In contrast, other formats such as the GFF family of formats, require *start ≤ end* regardless of the strand, which is equivalent to interpreting the INSDC location string as complement(*start*..*end*). This convention has some practical advantages when dealing with numerical operations on features sets, such as checking for overlaps or indexing data. For example, the feature length is given by *length* = *end - start* + 1 under this numerically convenient scheme where the interpretation of *start* versus *end* is strand independent.

There are a number of implicit conventions in INSDC data that need to be translated into the more explicit FALDO model. Some of the complicated regions for INSDC are features on a circular chromosome, the most common of which are features that overlap the chromosome’s origin of replication. One such feature is the “Protein II” gene from the reverse strand of f1 bacteriophage (ddbj:J02448). “Protein II” transcription starts at position 6006 on the reverse strand and ends at position 831 (see Figure 9).

**Figure 9:**
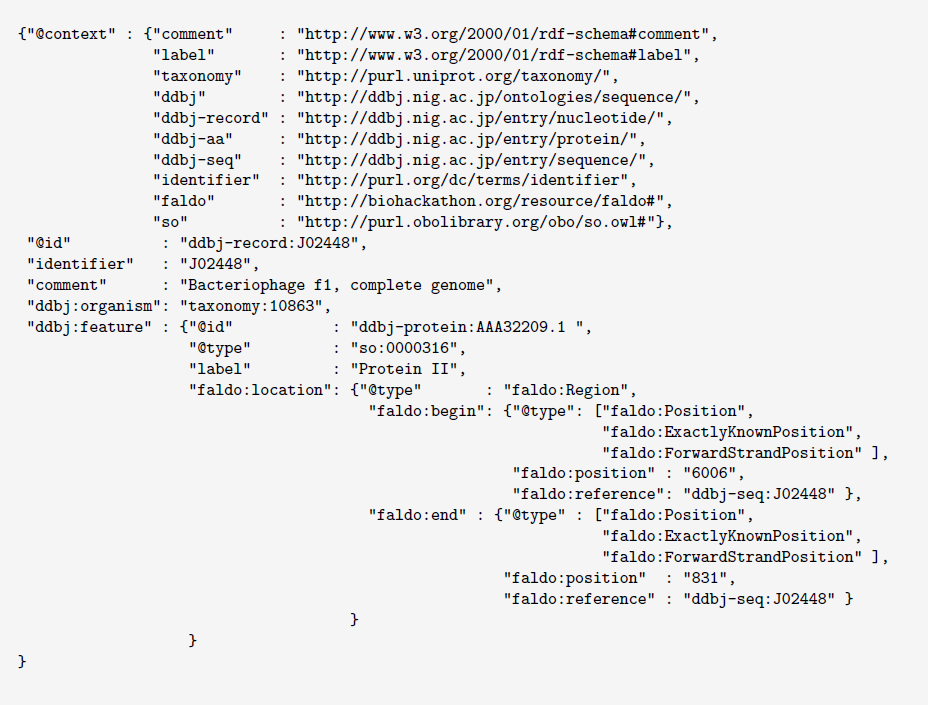
Partial example of using FALDO in JSON-LD [27] syntax to describe the CDS “Protein II” at join(6006..6407,1..831) on J02448.

### Fuzzy locations

Feature positions in, for example, INSDC or UniProt, are not always exactly known or described, but we should strive to describe our limited knowledge as accurately as possible. Take for example the position of the signal peptide annotation shown in Figure 6, where the protein sequence is known to belong to a family of proteins, but unfortunately only a part of the amino acid sequence is known. The UniProt curator deduced that the signal peptide region only partly overlaps the known sequence fragment. The same is true in the related INSDC record, were the CDS starts and ends before the known mRNA sequence (see Figure 7). As demonstrated in the figure, this limited knowledge can be described using the FALDO classes faldo: InRangePosition and faldo: OneOfPosition.

### Restriction enzymes

The task of describing the recognition sites of most restriction enzymes is quite straightforward, as is describing the cleavage site of a blunt end cutting enzyme. However, the cut site of a sticky-end cutting enzyme like HindIII that leaves an “overhang” is more challenging to specify, since it cuts in a different place on the forward and reverse strands. Figure 10 demonstrates how to describe this in FALDO by specifying start and end positions of the cut site that are on different strands.

**Figure 10:**
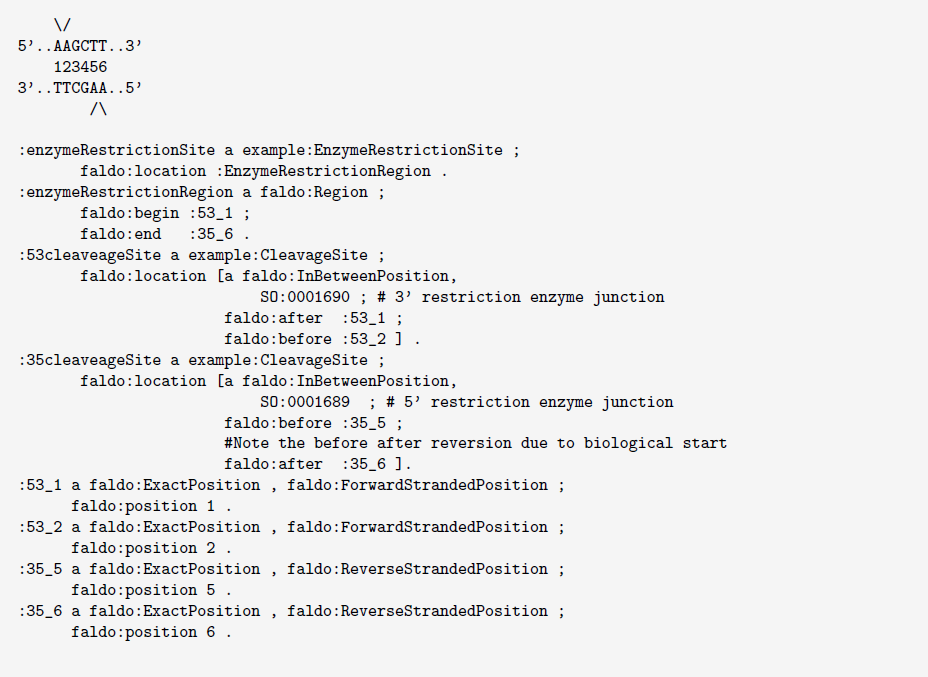
FALDO representation of HindIII restriction enzyme cleavage site with sticky ends.

## Discussion

When designing FALDO, a broad range of use cases were considered from human genome annotations to protein domains and glycan binding sites on amino acid sequences, with the goal of developing a scheme general enough to describe regions of DNA, RNA and protein sequences.

Advantages and drawbacks of existing file formats were considered, including line based column formats like BED and GTF/GFF3 which focus on exact ranges on a given sequence, and the more complex locations supported by the INSDC feature tables used by DDBJ, NCBI-GenBank and EMBL-Bank.

The simplest non-stranded range location on a linear sequence requires a start and end coordinate, but even here there are existing competing conventions for describing open or closed end-points using zero and one-based counting (for example BED versus GTF/GFF3/INSDC).

In FALDO we always count from the start of the forward 5’–3’ strand, even for features on the reverse strand. This encoding means there is no need to know the length of the sequence to compare positions on the different strands of a linear chromosome or genome. The end and start position of a region is inclusive. Unlike formats like GTF/GFF3, FALDO shares with Chado [28] the convention that the start coordinate should be the biological start (which may be a numerically higher value than the end coordinate).

For a semantic description describing the strand explicitly is preferable. FALDO chooses to add the strand information to the position. This is required to accurately describe for example the sticky ends of an enzyme digestion cut site, as in the HindIII example (Figure 10).

A major difference with other standards is that we chose to make strandedness and reference sequence a property of the position, instead of the region. This is important in a number of use cases. For example, one may need to describe the position of a gene on a draft genome assembly where the start and end are known to be on different contigs. This can be the case when RNA mapping is used in the genome assembly process. Another is when rough semantics are used in queries e.g. answering what is the start and end of a gene. In a process called transplicing, exons of one gene can be found on multiple chromosomes, or on different strands of the same chromosome. e.g. gene *mod* (*mdg4)* of *Drosophila melanogaster* (uniprot:Q86B87). In such cases the start of the gene can be on a different reference sequence or strand than the end. These biological realities cannot be described accurately if the reference sequence was a property of the region. As a side effect, it allows single nucleotide or amino acid sites to be described directly as a position without a need for an artificial region of length one.

Every faldo: Position refers to the sequence it is on. This allows us to say that gene *XX* starts at position 4 of assembly *Y* 1, while the same conceptual gene starts at position 5 of assembly *Y* 2. Even within the same assembly, FALDO offers the possibility to describe features in different contexts at the same time, allowing for instance to represent a SNP in terms of its position within a known coding region (i.e. gene coordinates) and within a chromosome region, which offers clear advantages for features annotation. Chado also allows multiple locations per feature, but unlike FALDO, the start and end of any location must be in the same region, which prohibits for example a feature that spans more than one contig, or describing the same feature on two different genome assemblies.

### Efficiency of Region-of-Interest queries

For FALDO we also considered query efficiency in comparison to existing search technology. Region of interest (ROI) queries are common operations performed on a set of genome annotations to extract a set of features within a range. For applications such as genome browsers, it is important that these are efficient enough. Although some RDF query engines may perform poorly when performing ROI queries over large feature sets, others have special indexes (e.g. literal filter indexes) that improve query performance. There is scope for further optimisation in the context of a SPARQL query by combining efficient algorithms and indexes such as Nested Containment Lists (NCLs) [29] or spatial indexes.

As a RDF based format, FALDO can be used to represent feature position information in a wide variety of serialisations e.g. JSON-LD, RDF/XML, Turtle, RDFa (embedded in HTML). This allows developers flexibility in consideration of their usage scenario, while at the same time allowing conversion to the common RDF triple model used in RDF databases and accessed by SPARQL queries.

## Conclusions

FALDO is a small ontology for describing biological features in a consistent manner that bioinformaticians can depend upon. The diverse software and high-profile databases already using FALDO show that it has enough power to describe existing biological feature locations. The uptake of this ontology means that it is now much easier for users querying biological databases on the Semantic Web to compare features on the basis of locations. This also means that visualisation tools that access positional data via SPARQL can easily reuse significant parts of queries between databases.

## Availability and requirements

FALDO is publicly available at the URL http://biohackathon.org/resource/faldo which is developed under source code control at https://github.com/JervenBolleman/FALDO hosted by GitHub Inc, where everyone is free to suggest extensions and improvements and if required extend FALDO to meet their unique requirements. FALDO currently uses the Creative Commons Attribution Zero 1.0 Public Domain dedication license, making FALDO available to use and reuse free of charge.

The ontology is shared in the Turtle (http://www.w3.org/TR/turtle/) RDF syntax, which can be automatically converted to another RDF syntax such as RDF/XML if required.

## List of abbreviations

BED: Browser Extensible Data (file format)
DDBJ: DNA Data Bank of Japan
EMBL: European Molecular Biology Laboratory
FALDO: Feature Annotation Location Description Ontology
GFF: Generic Feature Format
GFF3: Generic Feature Format version 3
GTF: Gene Transfer Format, a variant of GFF
GVF: Genome Variation Format, an extension to GFF3
INSDC: International Nucleotide Sequence Database Collaboration
OWL: Web Ontology Language (note acronym is OWL, not WOL)
RDF: Resource Description Framework
SPARQL: SPARQL Protocol and RDF Query Language
UniProtKB: Universal Protein Knowledgebase
VCF: Variant Call Format

## Authors contributions

Jerven Bolleman wrote the basic ontology file and mapping to UniProt RDF and contributed to the text of this article. Robert Hoehndorf did the GFF3 to OWL conversion. Peter Cock wrote a large section of this paper, and co-ordinated the working group during the BioHackathon meetings. Raoul JP Bonnal adapted BioRuby to match the ontology and wrote the Cufflinks to locations.rdf converter. Francesco Strozzi contributed to the Cufflinks RDF converter and wrote the VCF converter that uses FALDO for sequence variations positions. Takatomo Fujisawa and Toshiaki Katayama implemented an ontology and a tool for converting INSDC records to RDF and used FALDO to describe features on genomes in TogoGenome.

Robert Buels adapted JBrowse to query SPARQL endpoints that use this format to generate custom tracks. Joachim Baran incorporated FALDO into the genomic RDFization implementations of the BioInterchange software.

## Acknowledgments

The developers recognize the invaluable contributions from the community in helping to create this standard. We would like to especially thank the organisers and funders of the BioHackathon series of meetings for hosting the original discussions leading to FALDO (http://www.biohackathon.org/). The BioHackathon series, this work and Toshiaki Katayama were supported by National Bioscience Database Center (NBDC) of Japan Science and Technology Agency (JST) and Database Center for Life Science (DBCLS) of Research Organization of Information and Systems (ROIS) in Japan. Jerven Bolleman was supported in his role at Swiss-Prot, whose activities at the SIB Swiss Institute of Bioinformatics are supported by the Swiss Federal Government through the The State Secretariat for Education, Research and Innovation SERI. Christopher Mungall was supported by the NIH under R24OD011883 and by the Director, Office of Science, Office of Basic Energy Sciences, of the U.S. Department of Energy under Contract No. DE-AC02-05CH11231. Peter Cock was supported by the Scottish Government Rural and Environmental Research and Analysis Directorate. Takatomo Fujisawa was supported by the DNA Databank of Japan (DDBJ), Research Organization of Information and Systems (ROIS) in Japan.

